## Supplementary Tables S1-S5, Figures S1-S8, and Movie S1 legend for "Structural basis for saxitoxin congener binding and neutralization by anuran saxiphilins"

1 October 2024

[8]Molecular Biophysics and Integrated Bio-imaging Division
Lawrence Berkeley National Laboratory, Berkeley, CA 94720 USA

[2]Department of Chemistry
Stanford University, Stanford, CA 94305 USA

[3]National Oceanic and Atmospheric Administration
National Centers for Coastal Ocean Science
Charleston, SC 29412 USA

[4]Department of Pharmacological Sciences
Ichan School of Medicine at Mount Sinai
New York, NY 10029 USA

[#] Present address:
Department of Anatomy and Physiology
Shanghai Jiao Tong University School of Medicine
Shanghai, 200025, China

**Table S1 Thermofluor data comparison.**

| **Toxin** | ***Rc*Sxph ΔTm (°C)** | **ΔΔTm (°C)** | **n** | ***Np*Sxph ΔTm (°C)** | **ΔΔTm (°C)** | **n** |
|---|---|---|---|---|---|---|
| **STX** | 3.6 ± 0.2 | - | 8 | 3.2 ± 0.3 | - | 13 |
| **dcSTX** | 3.1 ± 0.1 | -0.5 ± 0.2 | 4 | 2.2 ± 0.1 | -1.0 ± 0.3 | 2 |
| **GTX2/3** | 2.7 ± 0.1 | -0.9 ± 0.2 | 4 | 2.2 ± 0.1 | -1.0 ± 0.3 | 4 |
| **dcGTX2/3** | 2.3 ± 0.1 | -1.3 ± 0.2 | 5 | 1.4 ± 0.1 | -1.8 ± 0.3 | 2 |
| **GTX5** | 3.7 ± 0.1 | 0.1 ± 0.2 | 4 | 3.3 ± 0.1 | 0.1 ± 0.3 | 2 |
| **C1/C2** | 2.6 ± 0.1 | -1.0 ± 0.2 | 4 | 1.5 ± 0.2 | -1.7 ± 0.4 | 4 |
| **neoSTX** | 0.7 ± 0.4 | -2.9 ± 0.4 | 6 | 0.3 ± 0.1 | -2.9 ± 0.3 | 4 |
| **dc-neoSTX** | 0.6 ± 0.3 | -3.0 ± 0.4 | 5 | 0.4 ± 0.2 | -2.8 ± 0.4 | 2 |
| **GTX1/4** | -0.2 ± 0.1 | -3.8 ± 0.2 | 4 | -0.1 ± 0.1 | -3.3 ± 0.3 | 3 |
| **GTX6** | 0.8 ± 0.3 | -2.8 ± 0.4 | 6 | 0.2 ± 0.2 | -3.0 ± 0.4 | 5 |

n, number of observations

$\Delta Tm = Tm_{Sxph+20\mu M\ toxin} - Tm_{Sxph}$

$\Delta\Delta Tm = Tm_{Sxph+20\mu M\ toxin} - Tm_{Sxph+20\mu M\ STX}$

ΔΔTm ranges are indicated as: White, STX; Green, ΔΔTm > -0.5 °C;
Yellow, -1.0 °C ≤ ΔΔTm ≤ -0.5 °C; Orange, -2.0 °C ≤ ΔΔTm < -1.0 °C; Red, ΔΔTm < -2.0 °C.

Errors are S.D.

**Table S2 Competition Fluorescence Polarization (FPc) data comparison.**

| Toxin | *Rc*Sxph Kd (nM) | ΔΔG (kcal mol$^{-1}$) | n | *Np*Sxph Kd (nM) | ΔΔG (kcal mol$^{-1}$) | n |
|---|---|---|---|---|---|---|
| **STX** | 8.9 ± 1.6 | - | 3 | 8.7 ± 1.7 | - | 4 |
| **dcSTX** | 20.3 ± 3.3 | 0.49 | 4 | 66 ± 16 | 1.19 | 4 |
| **GTX2/3** | 52.0 ± 8.3 | 1.04 | 3 | 133 ± 57 | 1.61 | 8 |
| **dcGTX2/3** | 180 ± 12 | 1.78 | 3 | 568 ± 68 | 2.47 | 3 |
| **GTX5** | 9.6 ± 0.8 | 0.04 | 3 | 7.3 ± 1.8 | -0.10 | 4 |
| **C1/C2** | 97.0 ± 8.3 | 1.41 | 3 | 186 ± 46 | 1.81 | 6 |
| **neoSTX** | >5000 | >3.7 | 3 | >5000 | >3.7 | 5 |
| **dc-neoSTX** | >5000 | >3.7 | 3 | >5000 | >3.7 | 6 |
| **GTX1/4** | >5000 | >3.7 | 3 | >5000 | >3.7 | 3 |
| **GTX6** | 1516 ± 222 | 3.04 | 3 | 1239 ± 578 | 2.94 | 4 |

n, number of observations

Kd, dissociation constant

$\Delta\Delta G = RT \ln(Kd_{toxin}/Kd_{STX})$  T = 298 K

ΔΔG ranges are indicated as: White, STX; Green, ΔΔG < 1.0 kcal mol$^{-1}$; Yellow, 1.0 ≤ ΔΔG < 2.0 kcal mol$^{-1}$; Orange, 2.0 ≤ ΔΔG ≤ 3.0 kcal mol$^{-1}$; Red, ΔΔG > 3.0 kcal mol$^{-1}$

Errors are S.D.

**Table S3 Radioligand receptor binding (RBA) data comparison.**

| Toxin | *Rc*Sxph Kd (nM) | ΔΔG (kcal mol$^{-1}$) | n | *Np*Sxph Kd (nM) | ΔΔG (kcal mol$^{-1}$) | n | Rat brain homogenate Kd (nM) | ΔΔG (kcal mol$^{-1}$) | n |
|---|---|---|---|---|---|---|---|---|---|
| **STX** | 7.1 ± 1.1 | - | 5 | 6.1 ± 1.3 | - | 7 | 1.5 ± 0.2 | - | 10 |
| **dcSTX** | 20 ± 4.0 | 0.61 | 5 | 40 ± 10 | 1.11 | 7 | 2.8 ± 0.3 | 0.37 | 2 |
| **GTX2/3** | 48 ± 10 | 1.13 | 5 | 52 ± 13 | 1.27 | 7 | 4.8 ± 1.1 | 0.69 | 4 |
| **dcGTX2/3** | 159 ± 6.0 | 1.84 | 5 | 396 ± 168 | 2.47 | 6 | 20 ± 4.0 | 1.53 | 4 |
| **GTX5** | 5.0 ± 1.6 | -0.21 | 6 | 3.2 ± 2.3 | -0.38 | 5 | 326 ± 72 | 3.19 | 3 |
| **C1/C2** | 106 ± 7.0 | 1.60 | 3 | 255 ± 16 | 2.21 | 3 | 221 ± 91 | 2.96 | 3 |
| **neoSTX** | >1000 | >3.0 | 2 | >1000 | >3.0 | 3 | 0.5 ± 0.1 | -0.65 | 2 |
| **dc-neoSTX** | >1000 | >3.0 | 2 | >1000 | >3.0 | 4 | 61 ± 39 | 2.19 | 4 |
| **GTX1/4** | >1000 | >3.0 | 2 | >1000 | >3.0 | 3 | 3.2 ± 1.0 | 0.45 | 4 |
| **GTX6** | >1000 | >3.0 | 5 | >1000 | >3.0 | 7 | 88 ± 5.1 | 2.41 | 3 |

n, number of observations

$\Delta\Delta G = RT \ln(Kd_{toxin}/Kd_{STX})$  T = 298 K

ΔΔG ranges are indicated as: White, STX; Green, ΔΔG < 1.0 kcal mol$^{-1}$; Yellow, 1.0 ≤ ΔΔG < 2.0 kcal mol$^{-1}$; Orange, 2.0 ≤ ΔΔG ≤ 3.0 kcal mol$^{-1}$; Red, ΔΔG > 3.0 kcal mol$^{-1}$

Errors are S.D.

**Table S4 Crystallographic data collection and refinement statistics**

| | *Np*Sxph:dcSTX (co-crystal) PDB: 8V68 | *Np*Sxph:GTX2 (co-crystal) PDB: 8V69 | *Np*Sxph:dcGTX2 (co-crystal) PDB: 8V65 | *Np*Sxph:GTX5 (co-crystal) PDB: 8V66 | *Np*Sxph:C1 (co-crystal) PDB: 8V67 |
|---|---|---|---|---|---|
| **Data Collection** | | | | | |
| Space group | R3 | R3 | R3 | R3 | R3 |
| Cell dimensions a/b/c (Å) | 229.47, 229.47, 67.61 | 229.74, 229.74, 67.87 | 229.67, 229.67, 67.71 | 229.16, 229.16, 67.50 | 229.78, 229.78, 67.39 |
| α/β/γ (°) | 90, 90, 120 | 90, 90, 120 | 90, 90, 120 | 90, 90, 120 | 90, 90, 120 |
| Resolution (Å) | 43.37-1.9 (1.968-1.9) | 43.42-1.95 (2.02-1.95) | 43.4-1.8 (1.864-1.8) | 43.31-1.9 (1.968-1.9) | 43.43-1.90 (1.968-1.90) |
| Rmerge (%) | 0.1038 (3.545) | 0.09641 (2.338) | 0.1286 (5.018) | 0.1362 (2.498) | 0.2557 (3.836) |
| I / σ I | 12.65 (1.03) | 9.91 (0.81) | 9.74 (0.66) | 8.15 (1.19) | 6.51 (0.69) |
| CC(1/2) | 0.999 (0.49) | 0.999 (0.41) | 0.998 (0.394) | 0.998 (0.58) | 0.997 (0.356) |
| Completeness (%) | 99.57 (95.89) | 99.98 (100.00) | 99.97 (99.90) | 99.98 (99.99) | 99.92 (99.50) |
| Redundancy | 15.1 (15.5) | 10.0 (10.2) | 14.9 (14.7) | 15.1 (15.6) | 30.2 (31.0) |
| Total reflections | 1583855 (161838) | 972728 (98797) | 1843446 (181645) | 1577994 (162070) | 3161523 (322475) |
| Unique reflections | 104618 (10454) | 97333 (9717) | 123415 (12335) | 104167 (10420) | 104543 (10419) |
| Wilson B-factor | 43.54 | 49.43 | 43.13 | 42.78 | 44.84 |
| Wavelength (Å) | 1.116 | 1.116 | 1.116 | 1.116 | 1.116 |
| **Refinement** | | | | | |
| $R_{work}$ / $R_{free}$ (%) | 19.65/22.43 | 18.68/21.16 | 19.88/21.82 | 19.00/21.82 | 20.70/22.96 |
| No. of chains in AU | 1 | 1 | 1 | 1 | 1 |
| No. of protein atoms | 6392 | 6381 | 6373 | 6402 | 6382 |
| No. of ligand atoms | 34 | 42 | 16 | 41 | 46 |
| No. of water atoms | 400 | 273 | 428 | 378 | 291 |
| RMSD bond lengths (Å) | 0.003 | 0.014 | 0.004 | 0.007 | 0.009 |
| RMSD angles (°) | 0.57 | 1.24 | 0.68 | 0.90 | 1.03 |
| Ramachandran favored/allowed/outliers (%) | 96.95/2.93/0.12 | 95.96/3.92/0.12 | 96.93/2.82/0.25 | 96.21/3.54/0.24 | 95.95/3.93/0.12 |

**Table S5 *Hs*Na$_V$1.4 and *Pt*Na$_V$1.4 toxin responses.**

| Toxin | *Hs*Na$_V$1.4 | | | *Pt*Na$_V$1.4 | | |
|---|---|---|---|---|---|---|
| | $IC_{50}$ (nM) | $IC_{90}$ (nM) | n | $IC_{50}$ (nM) | $IC_{90}$ (nM) | n |
| **STX** | 3.0 ± 1.6 | 50 | 9 | 12.6 ± 1.4* | 100 | 6 |
| **dcSTX** | 14.7 ± 8.7 | 200 | 4 | 144.4 ± 29.7 | 800 | 6 |
| **GTX2/3** | 6.8 ± 1.1 | 60 | 8 | 27.4 ± 2.1 | 200 | 6 |
| **dcGTX2/3** | 31.8 ± 10.6 | 300 | 6 | N/A | N/A | N/A |
| **GTX5** | 335.3 ± 55.4 | 2700 | 8 | 2290 ± 776 | >10,000 | 6 |
| **C1/C2** | 151.5 ± 37.0 | 1000 | 5 | N/A | N/A | N/A |

$IC_{50}$, half-maximal inhibitory concentration

$IC_{90}$, concentration required to block 90% of the current

n, number of cells

Errors are S.D.

*data taken from [1]

**Figure S1** **Zakrzewska *et al.***

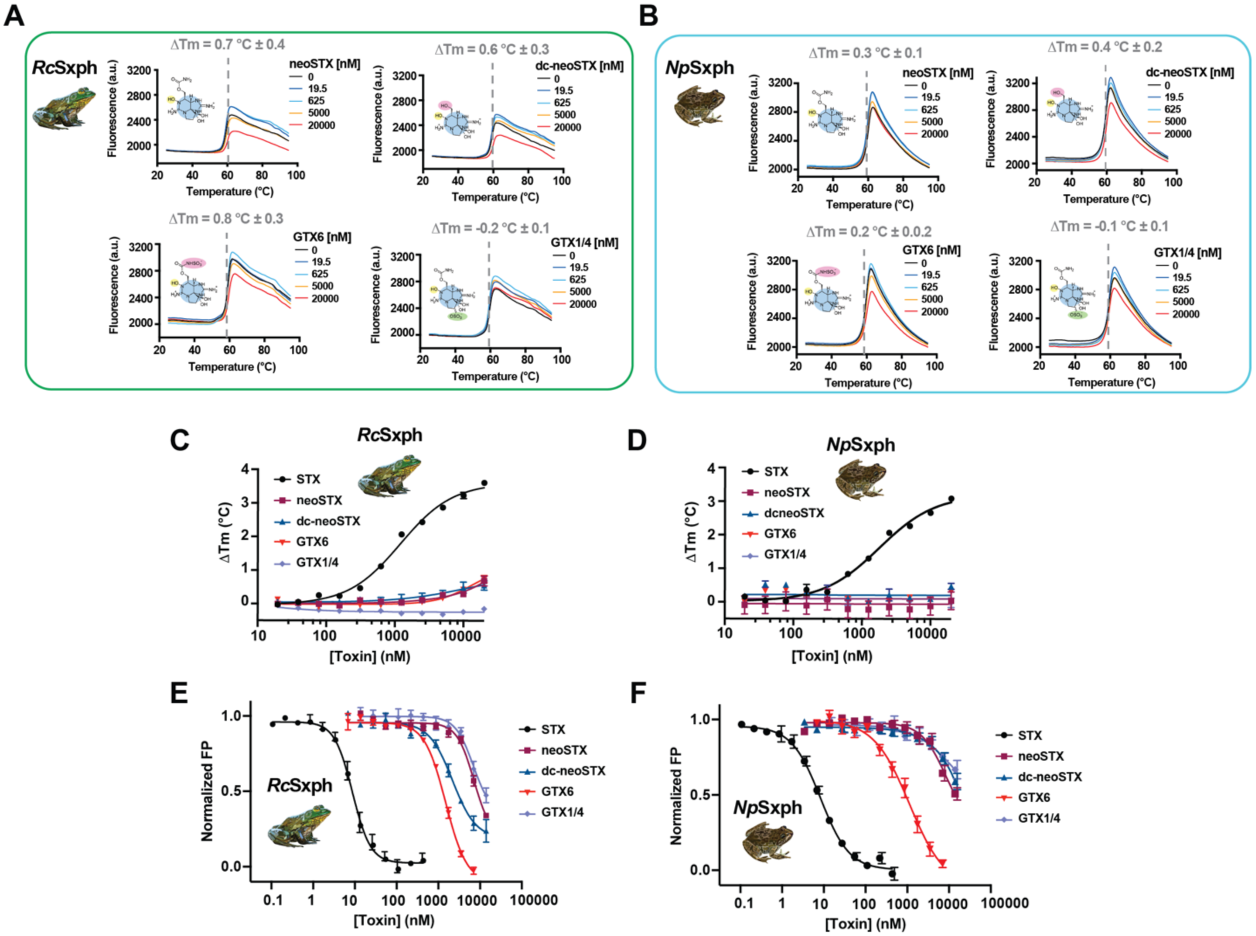


**Figure S1 *Rc*Sxph and *Np*Sxph poorly interact with neo-STX congeners. A,** and **B,** Exemplar TF assay results for **A,** *Rc*Sxph and **B,** *Np*Sxph in the presence of the indicated concentrations of neoSTX, dc-neoSTX, GTX6, and GTX1/4. Toxin concentrations are 0 nM (black), 19.5 nM (blue), 625 nM (cyan), 5,000 nM (orange), and 20,000 nM (red). **C,** and **D,** Temperature dependence of ΔTm as a function of toxin concentration for TF assays of **C,** *Rc*Sxph and **D,** *Np*Sxph with STX (black circles), neoSTX (maroon squares), dc-neoSTX (blue triangles), GTX6 (red inverted triangles), and GTX1/4 (lavender diamonds). **E,** and **F,** FP competition binding assays for **E,** *Rc*Sxph and **F,** *Np*Sxph with STX (black circles), neoSTX (maroon squares), dc-neoSTX (blue triangles), GTX6 (red inverted triangles), and GTX1/4 (lavender diamonds). Error bars are S.E.M.

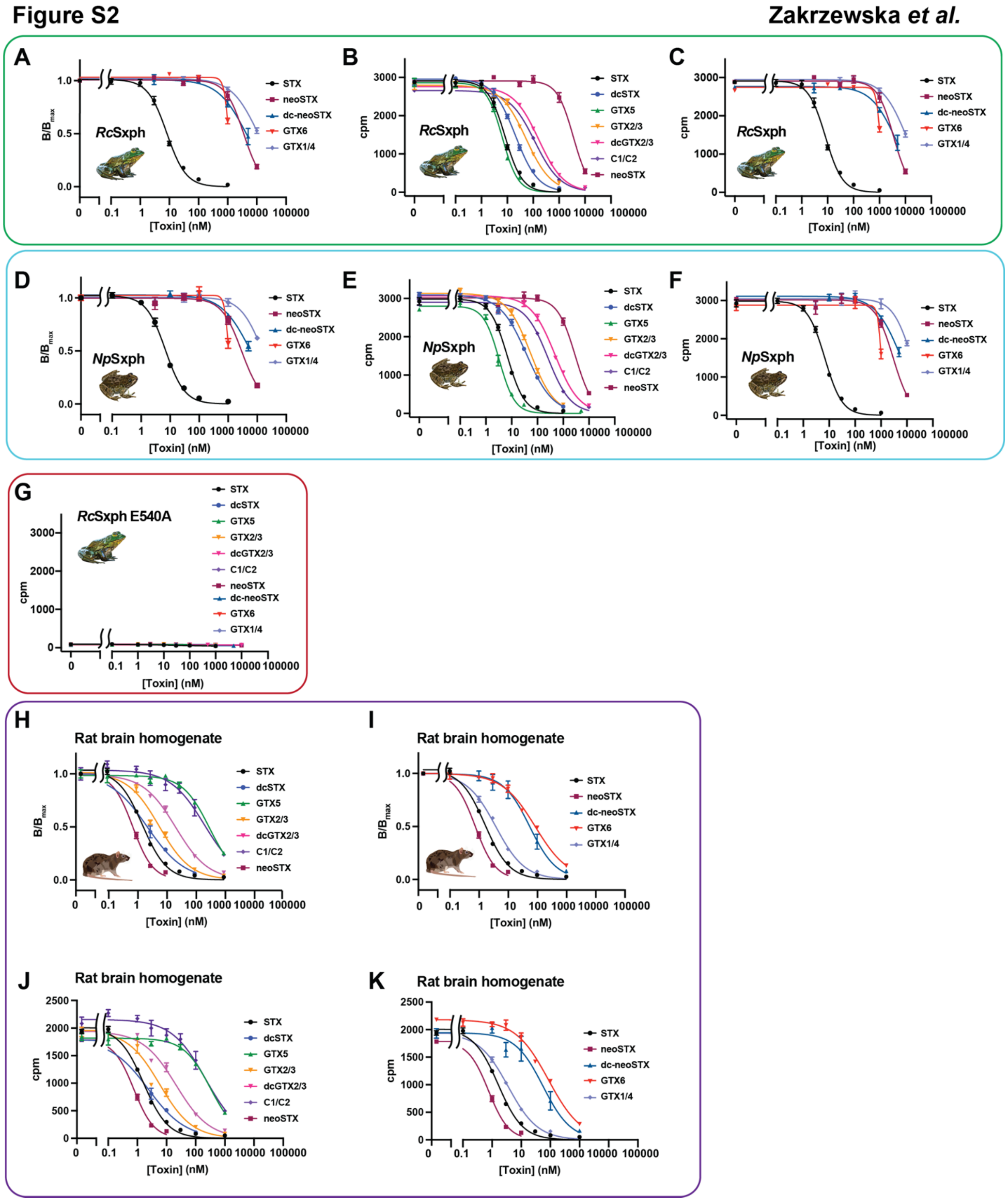


**Figure S2 Receptor binding assay comparisons. A,** *Rc*Sxph B/$B_{max}$ normalized RBA data for the indicated toxin, where B represents the bound [$^3$H]STX in the sample and $B_{max}$ is the maximum binding of [$^3$H]STX in the absence of competing unlabeled toxin. **B,** and **C**, Raw radioactive counts per minute (cpm) for *Rc*Sxph RBAs. **D,** *Np*Sxph normalized RBA data for the indicated toxins. **E,**

and **F**, Raw cpm for *Np*Sxph RBAs. **G,** Raw cpm for *Rc*Sxph E540A RBA. **H,** and **I,** Normalized RBA data for rat brain homogenate binding of **H,** STX, STX congeners, and neo-STX and **I,** STX, neo-STX, and neo-STX congeners. **J,** and **K**, Raw cpm for rat brain homogenate RBAs. In all panels toxins indicated as STX (black circles), dcSTX (blue circles), GTX5 (green triangles), GTX2/3 (orange inverted triangles), dcGTX2/3 (magenta inverted triangles), C1/C2 (lavender diamonds), neoSTX (dark red squares), dc-neoSTX (blue triangles), GTX6 (inverted red triangles), and GTX1/4 (light purple diamonds). Error bars are S.E.M..

**Figure S3** **Zakrzewska *et al.***

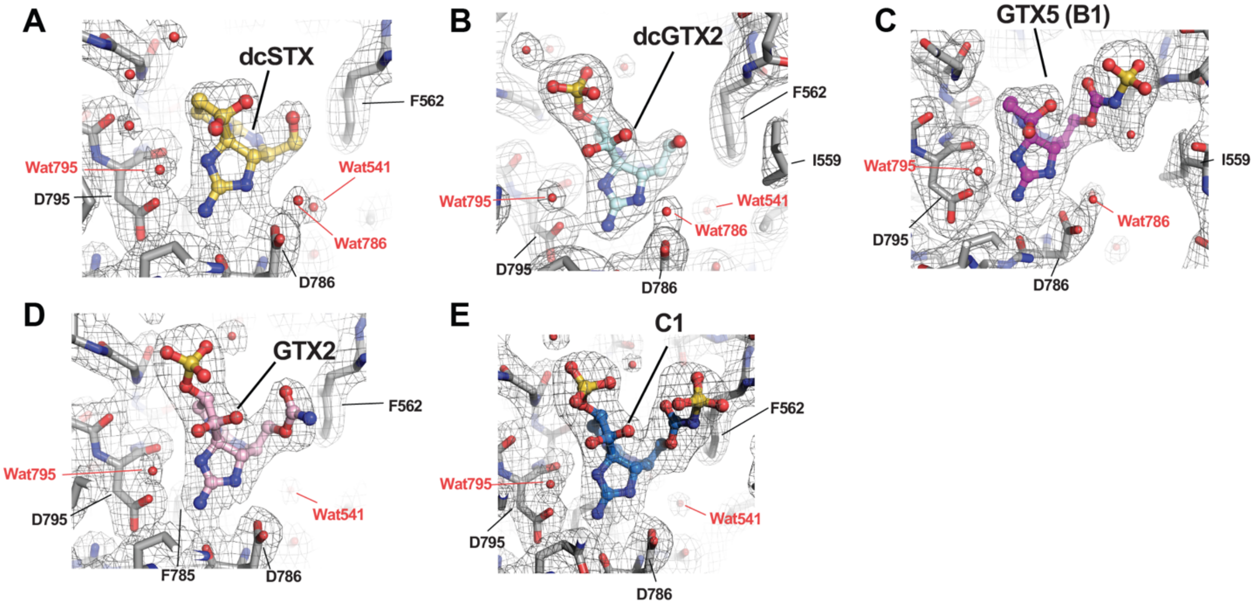


**Figure S3 Structures of *Np*Sxph:STX congener complexes. A-E,** Exemplar electron density (1σ) for **A,** *Np*Sxph:dcSTX (yellow). **B,** *Np*Sxph:dcGTX2 (cyan). **C,** *Np*Sxph:GTX5 (B1) (magenta). **D,** *Np*Sxph:GTX2 (light pink). **E,** *Np*Sxph:C1 (marine). *Np*Sxph is light grey. Water molecules (red) are shown as spheres. Select residues are labeled.

### Figure S4

Zakrzewska *et al.*

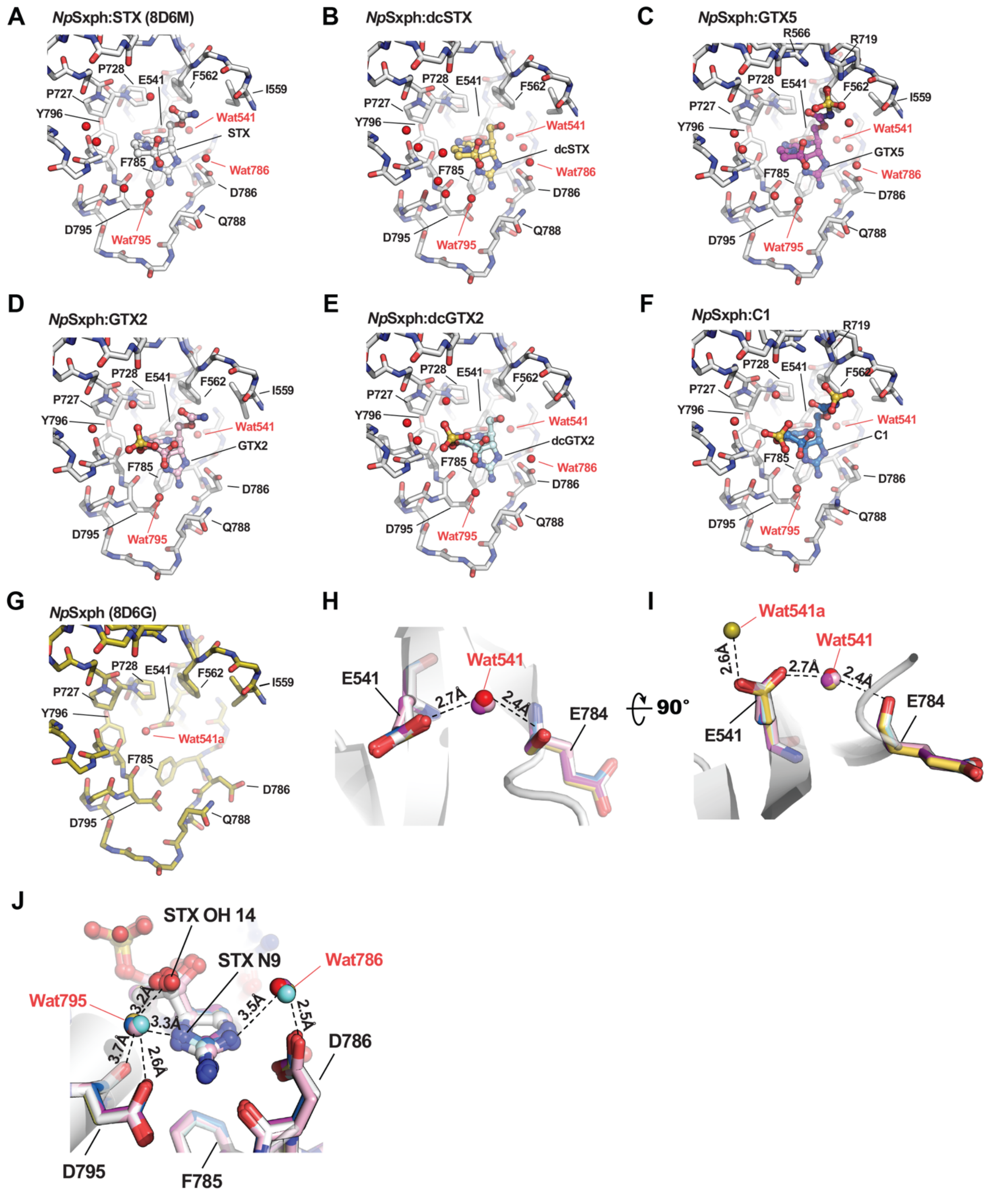


**Figure S4 *Np*Sxph STX binding pocket water network structure. A-F,** *Np*Sxph:toxin complexes (white) showing the positions of crystallographic water molecules (red spheres) in the toxin binding pocket for **A,** *Np*Sxph:STX (white) (PDB:8D6M) [2]. **B,** *Np*Sxph:dcSTX (yellow). **C,** *Np*Sxph:GTX5 (magenta). **D,** *Np*Sxph:GTX2 (light pink). **E,** *Np*Sxph:dcGTX2 (cyan). **F,** *Np*Sxph:C1 (marine). **G,** apo-*Np*Sxph (PDB:8D6G) (yellow orange) [2]. **H,** Close up view of Wat541 and coordinating residues Glu541 and Glu784 for *Np*Sxph:STX (white), *Np*Sxph:dcSTX (yellow), *Np*Sxph:dcGTX2 (cyan), *Np*Sxph:GTX5 (magenta), *Np*Sxph:GTX2 (light pink) and, *Np*Sxph:C1 (marine). **I,** Close up view of the positions of Wat541a and Wat541. Colors are as in 'H'. apo-*Np*Sxph (PDB:8D6G) (yellow orange). **J,** Close up view of the positions of Wat786 and Wat795 for *Np*Sxph:STX (white), *Np*Sxph:dcSTX (yellow), *Np*Sxph:dcGTX2 (cyan), *Np*Sxph:GTX5 (magenta), *Np*Sxph:GTX2 (light pink) and, *Np*Sxph:C1 (marine). STX positions of N9 and OH 14 are indicated.

**Figure S5** **Zakrzewska *et al.***

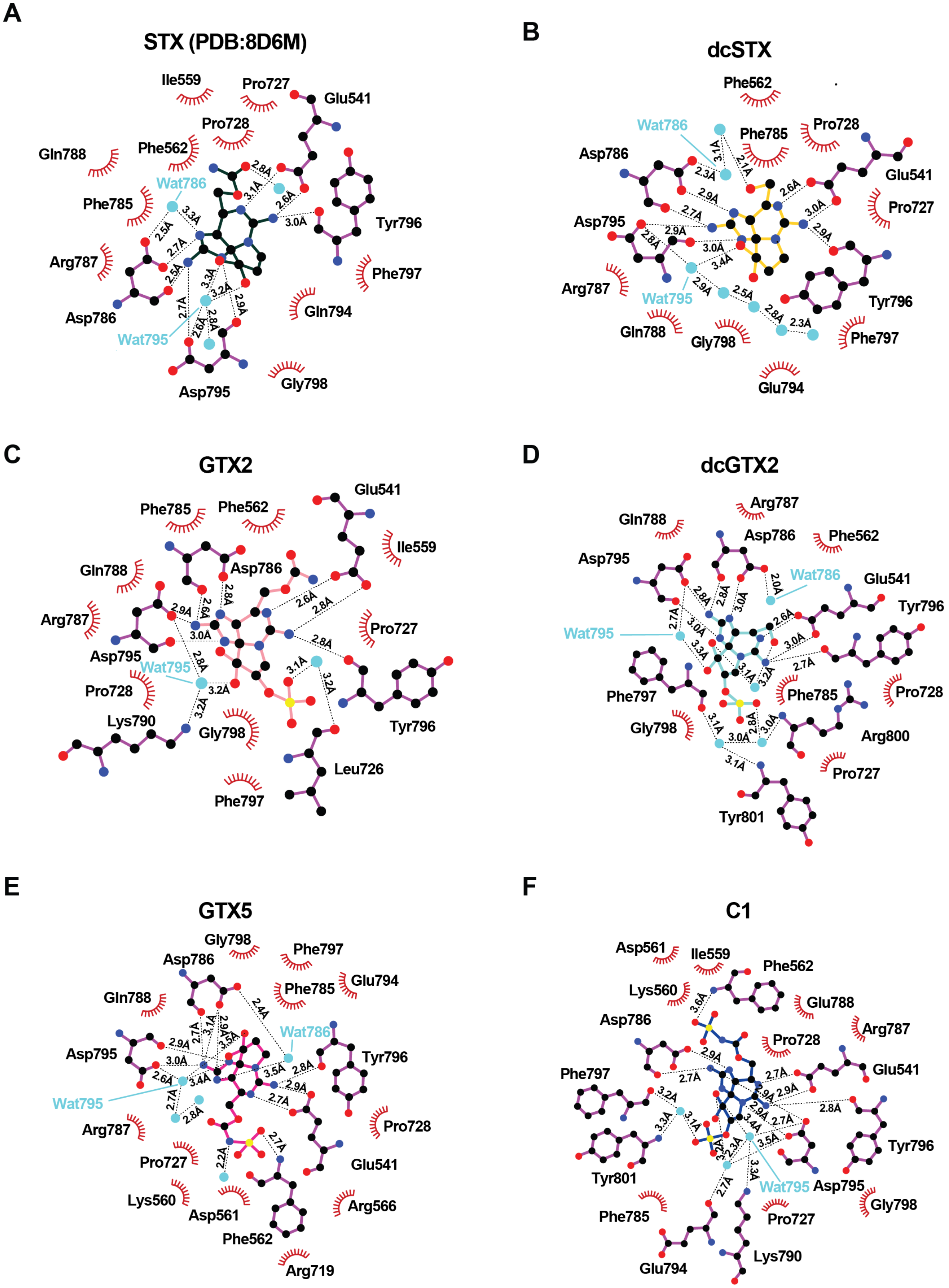

**Figure S5 *Np*Sxph:STX congener interactions. A-F,** LIGPLOT [3] diagrams showing interactions (5.0Å cutoff) for: **A**, *Np*Sxph:STX (PDB:8D6M) [2]. **B,** *Np*Sxph:dcSTX (yellow). **C,** *Np*Sxph:GTX2 (light pink). **D,** *Np*Sxph:dcGTX2 (cyan). **E,** *Np*Sxph:GTX5 (magenta). **F,** *Np*Sxph:C1 (marine).

**Figure S6** **Zakrzewska *et al.***

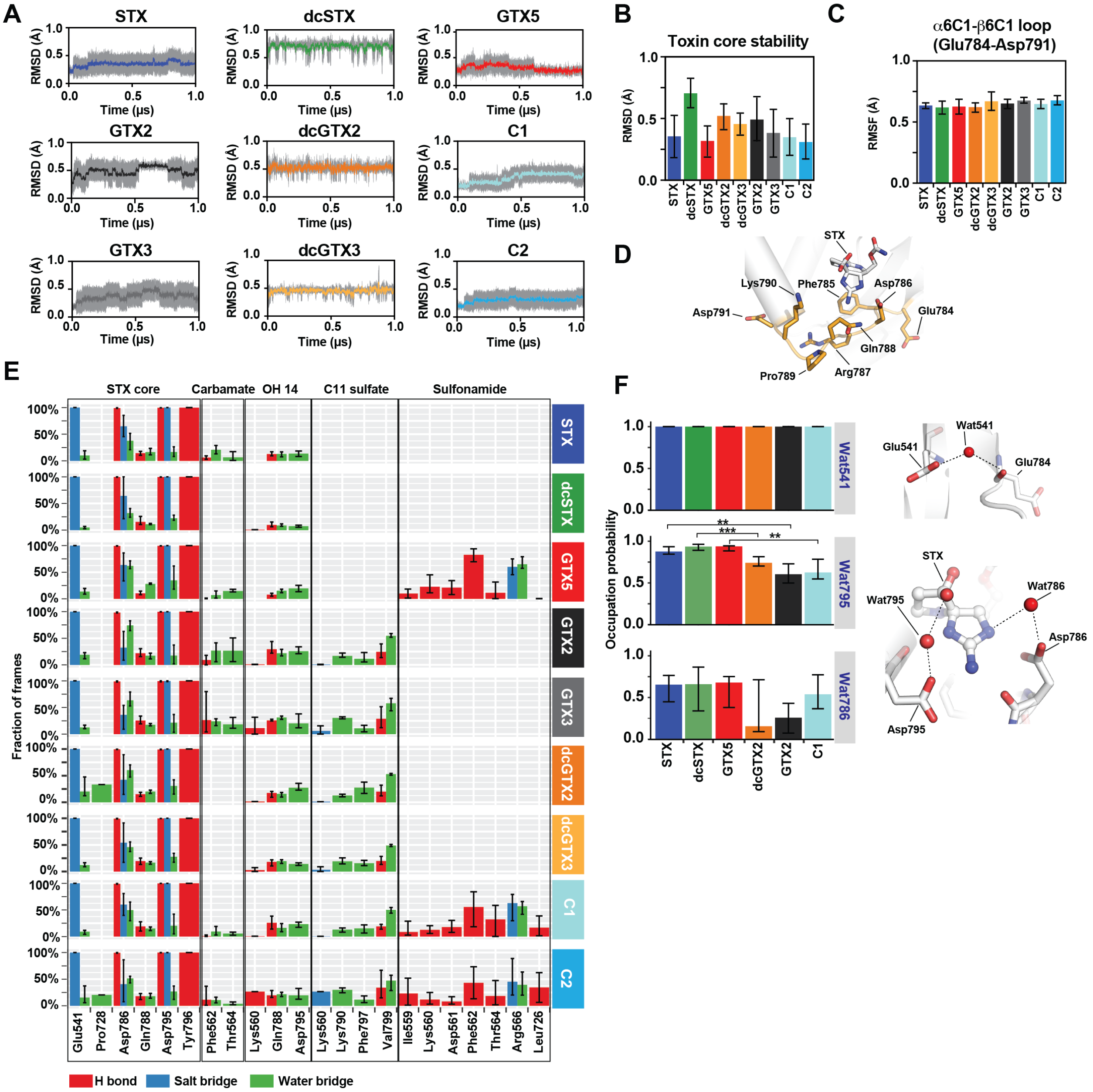


**Figure S6 *Np*Sxph:toxin simulations highlight ligand stability and effects on water network.** **A,** Time series of RMSD calculated from the crystallographic pose for non-hydrogen ligand atoms common to all STX congeners during 1 µs simulations (n=5). **B,** Mean and standard deviation of the average RMSD for the STX core atoms over the entire trajectory. **C,** Mean and standard deviation of the average RMSF for *Np*Sxph α6C1-β6C1 loop residues Cα atoms during 1 µs simulations (n=5). **D,** *Np*Sxph α6C1-β6C1 loop (orange) position in the *Np*Sxph:STX complex (PDB 8D6M)[2]. **E,** Structural interaction fingerprint (SIFt) analyses of *Np*Sxph complexes grouped by interaction type, protein residue, and interacting toxin moiety. **F,** (left) Mean probability of occupancy of the indicated hydration sites (Wat541, Wat795, and Wat786) for all simulated systems. For Wat795, a Welch two-sample two-sided t-test reveals significant differences in mean occupancy between dcGTX2 and dcSTX, GTX2 and STX, and GTX5 and C1, with p-values of 0.00219, 0.0137, and 0.0129, respectively. For Wat786, only the comparison between GTX2 and STX shows a significant, albeit weak, difference, (p-value = 0.0234). Comparisons between dcGTX2 and dcSTX and between GTX5 and C1 yield p-values > 0.1. (right) Locations of Wat541, Wat795, and Wat786 in the *Np*Sxph:STX structure (PDB: 8D6M)[2]. Dashed lines indicate hydrogen bonds. Error bars represent S.D. of the mean fraction of frames over five trajectories. ** $p < 0.02$, *** $p < 0.005$

**Figure S7** **Zakrzewska *et al.***

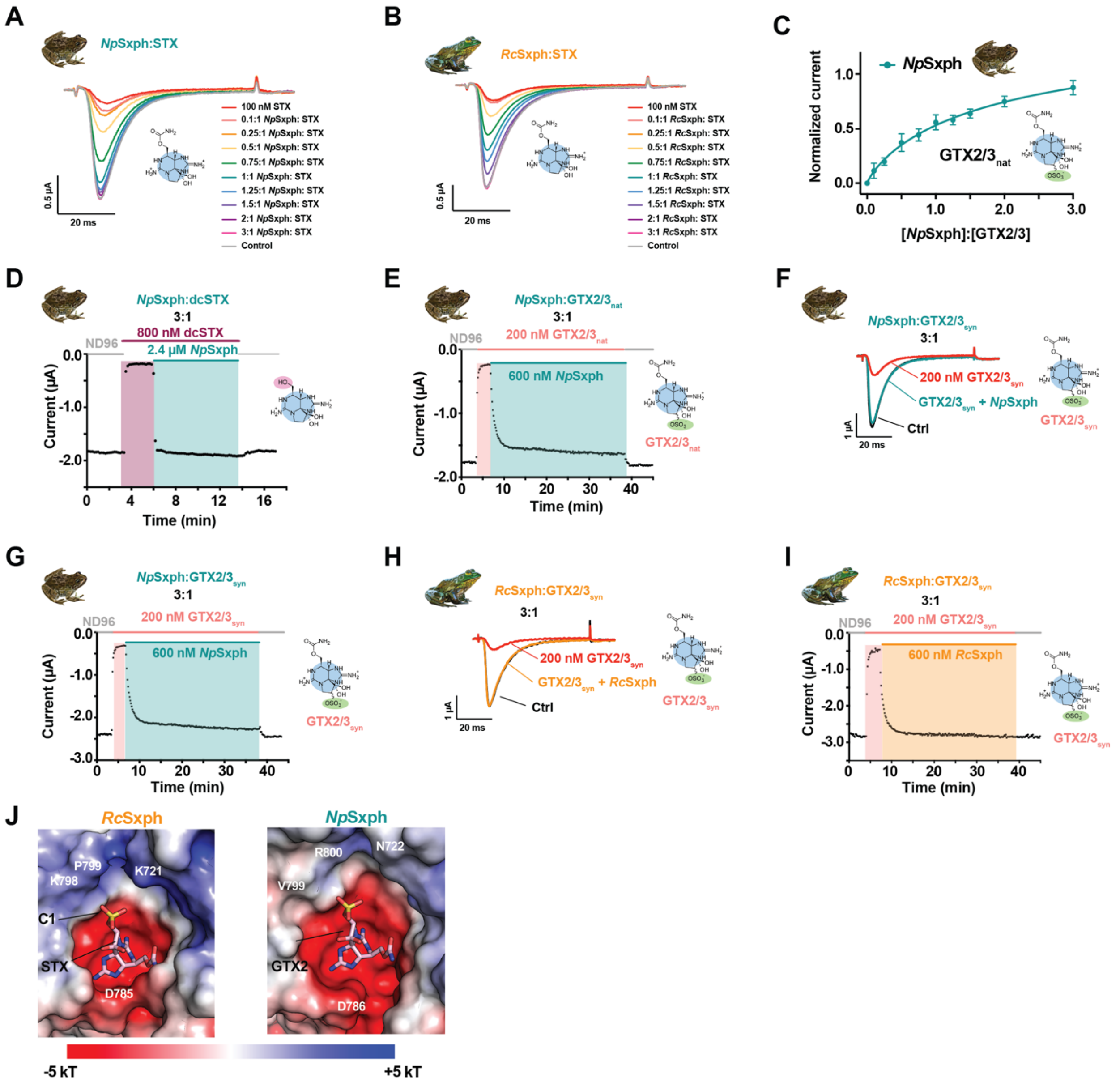


**Figure S7 Sxphs rescue *Pt*Na$_V$1.4 from toxin block. A** and **B,** Exemplar two electrode voltage clamp (TEVC) recordings of *Pt*Na$_V$1.4 expressed in *Xenopus* oocytes in the presence of 100 nM STX, the indicated Sxph:STX ratios, or control conditions for **A,** *Np*Sxph and **B,** *Rc*Sxph. **C,** [*Np*Sxph]:[GTX2/3] dose-response curve in the presence of 200 nM GTX2/3. **D,** and **E,** *Pt*Na$_V$1.4 responses to application of **D,** 800 nM dcSTX (purple) and **E,** 200 nM GTX2/3$_{nat}$ (light red) and 3:1 *Np*Sxph:toxin application (bluegreen). **F,** Exemplar TEVC recordings of *Pt*Na$_V$1.4 expressed in *Xenopus* oocytes in the presence of 200 nM GTX2/3$_{syn}$ and 3:1 *Np*Sxph:GTX2/3$_{syn}$. **G,** *Pt*Na$_V$1.4 response to application of 200 nM GTX2/3$_{syn}$ (light red) and 3:1 *Np*Sxph:toxin application (bluegreen). **H,** Exemplar TEVC recordings of *Pt*Na$_V$1.4 expressed in *Xenopus*

oocytes in the presence of 200 nM GTX2/$3_{syn}$ and 3:1 *Rc*Sxph:GTX2/$3_{syn}$. **I,** *Pt*$Na_V$1.4 response to application of 200 nM GTX2/$3_{syn}$ (light red) and 3:1 *Rc*Sxph:toxin application (orange). **J,** Comparisons of the electrostatic surface potentials of the toxin binding pocket from *Rc*Sxph:STX (PDB:6O0F)[4] with *Np*Sxph:GTX2 (PDB:8V69)[2]. STX and GTX2 are shown as sticks and space filling.

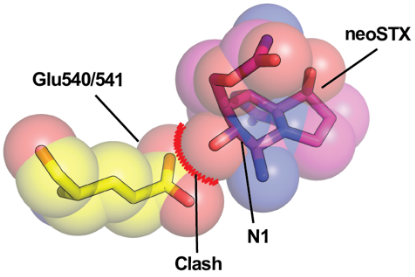


**Figure S8 Model of Sxph STX binding pocket:neoSTX clash.** Superposition of neoSTX on STX in the *Rc*Sxph:STX complex (PDB:6O0F)[4] indicates clash of the neoSTX N1 hydroxyl with the conserved glutamate (*Rc*Sxph540, *Np*Sxph541).

**Movie S1 Conformational changes between *Np*Sxph:STX congener structures.** Morph between the apo-*Np*Sxph (PDB:8D6G)[2] and each *Np*Sxph:STX congener structures showing the toxin binding site.

### References

1. Abderemane-Ali, F. et al. Evidence that toxin resistance in poison birds and frogs is not rooted in sodium channel mutations and may rely on "toxin sponge" proteins. *J Gen Physiol* **153**(2021).
2. Chen, Z. et al. Definition of a saxitoxin (STX) binding code enables discovery and characterization of the anuran saxiphilin family. *Proc Natl Acad Sci U S A* **119**, e2210114119 (2022).
3. Wallace, A.C., Laskowski, R.A. & Thornton, J.M. LIGPLOT: a program to generate schematic diagrams of protein-ligand interactions. *Protein Eng* **8**, 127-34 (1995).
4. Yen, T.-J., Lolicato, M., Thomas-Tran, R., Du Bois, J. & Minor, D.L., Jr. Structure of the Saxiphilin:saxitoxin (STX) complex reveals a convergent molecular recognition strategy for paralytic toxins. *Sci Adv* **5**(2019).
